# Methodological considerations on selection of stable reference genes for RT-qPCR in the neonatal rat brain in hypoxia and hypothermia

**DOI:** 10.1101/793786

**Authors:** M. Bustelo, M.A. Bruno, C.F. Loidl, H.W.M. Steinbusch, A.W.D. Gavilanes, D.L.A. van den Hove

## Abstract

Real-time reverse transcription PCR (qPCR) normalized to an internal reference gene (RG), is a frequently used method for quantifying gene expression changes in neuroscience. Although RG expression is assumed to be constantly independent of physiological or experimental conditions, several studies have shown that commonly used RGs are not expressed stably. The use of unstable RGs has a profound effect on the conclusions drawn from studies on gene expression, and almost universally results in spurious estimation of target gene expression. Approaches aimed at selecting and validating RGs often make use of different statistical methods, which may lead to conflicting results. The present study evaluates the expression of 5 candidate RGs (*Actb*, *Pgk1*, *Sdha*, *Gapdh*, *Rnu6b*) as a function of hypoxia exposure and hypothermic treatment in the neonatal rat cerebral cortex –in order to identify RGs that are stably expressed under these experimental conditions– and compares several statistical approaches that have been proposed to validate RGs. In doing so, we first analyzed the RG ranking stability proposed by several widely used statistical methods and related tools, i.e. the Coefficient of Variation (CV) analysis, GeNorm, NormFinder, BestKeeper, and the ΔCt method. Subsequently, we compared RG expression patterns between the various experimental groups. We found that these statistical methods, next to producing different rankings per se, all ranked RGs displaying significant differences in expression levels between groups as the most stable RG. As a consequence, when assessing the impact of RG selection on target gene expression quantification, substantial differences in target gene expression profiles were observed. As such, by assessing mRNA expression profiles within the neonatal rat brain cortex in hypoxia and hypothermia as a showcase, this study underlines the importance of further validating RGs for each new experimental paradigm considering the limitations of each selection method.

## Introduction

In qPCR analysis, reference genes (RGs) with stable expression levels are essential internal controls for relative quantification of mRNA expression. RGs normalize variations of candidate gene expression under different conditions (1,2). The ideal RG should be expressed at constant levels regardless of e.g. experimental conditions, developmental stages or treatments (3,4), and should have expression levels comparable to that of the target gene (5). Nevertheless, increasing evidence suggests that the expression of commonly used RGs often varies considerably under different experimental conditions, as reviewed previously (6,7). The choice of unstable RGs for the normalization of qPCR data may give rise to inaccurate results, concomitant with potential expression changes in genes of interest being easily missed or overemphasized. Thus, the identification of stable RGs is a prerequisite for reliable qPCR experiments (9–11).

RG selection should be performed using the same samples that will be compared when looking at genes of interest. For this purpose, several statistical methods have been proposed, i.e. GeNorm (12), qBase (13), BestKeeper (14), NormFinder (15), Coefficient of Variation (CV) analysis (16), and the comparative ΔCt method (17). These statistical methods rank the stability of the candidate RGs based on a unique set of assumptions and associated algorithms. As a result, the predictions of these methods can differ significantly based on the method used, potentially leading to conflicting results. This observation has been frequently made, but still seems to be systematically ignored in recent validation studies.

To address this issue, several approaches that make use of several statistical approaches at the same time, have been proposed, including i) a weighted rank (18–20), an approach that is compromised by the fact that it does not consider the strengths and drawbacks of each method for a given experimental setting; ii) the “Geometric mean rank” that uses the average of the stability ranks across different methods yielding an overall ranking (12,21); as well as iii) an integrated approach (22), including a first selection step making use of the CV analysis (eliminating genes with CV>50%), and subsequently ranking the remaining genes using GeNorm.

In the present study, we compared these methods, on the evaluation of the stability of 5 candidate RGs in a murine model of perinatal asphyxia and therapeutic hypothermia. Perinatal asphyxia is a clinical condition defined as oxygen deprivation that occurs around the time of birth and may be caused by perinatal events such as placental abruption, cord-prolapse, or tight nuchal cord, limiting the supply of oxygenated blood to the fetus (23). Recently, hypothermia has emerged as the standard of care for perinatal asphyxia. Although this treatment has been demonstrated to be effective in reducing mortality and long-term consequences of perinatal asphyxia, the underlying mechanisms of this therapy are still not completely understood (24–28). Assessing gene expression changes in the neonatal hypoxic-ischemic brain may be of added value in order to further decipher the mechanism of perinatal asphyxia and to increase the effectivity of therapeutic hypothermia and related therapies. Here, we used a murine perinatal hypoxic-ischemic encephalopathy model (29–31) to address the abovementioned problems in RG selection and qPCR normalization. Several *in vivo* and *in vitro* studies on hypoxia, making use of qPCR, have been reported (19,32–39), indicating that hypoxia significantly impacts the expression of various commonly used RGs. Although some of these studies use the same or similar hypoxia models, the results vary substantially across studies, emphasizing the need to publish these validation studies prior or parallel to reporting qPCR results.

We selected five candidate RGs based on published RG validation studies involving hypoxia (Table 1). Subsequently, we applied various validated methodological and statistical methods to evaluate the effects of anoxia and hypothermia on the expression stability of the candidate RGs. To evaluate the impact of the resulting RG selection, we assessed the expression levels of the Repressor Element 1-Silencing Transcription Factor (*Rest*), a gene that has been shown to be upregulated by hypoxic-ischemic injury in the peri-infarct cortex of adult rats following transient focal ischemia induced by middle cerebral artery occlusion (MCAO) (40). Moreover, the proapoptotic gene BCL2/BCL-XL-associated death promoter (*Bad*), a gene that has been shown to be up-regulated by hypoxia in the MCAO rat model, was assessed (41). This study provides a basis for the selection of RGs and useful guidelines for future gene expression studies, in particular regarding studies on developmental hypoxic insults.

**Table 1.**
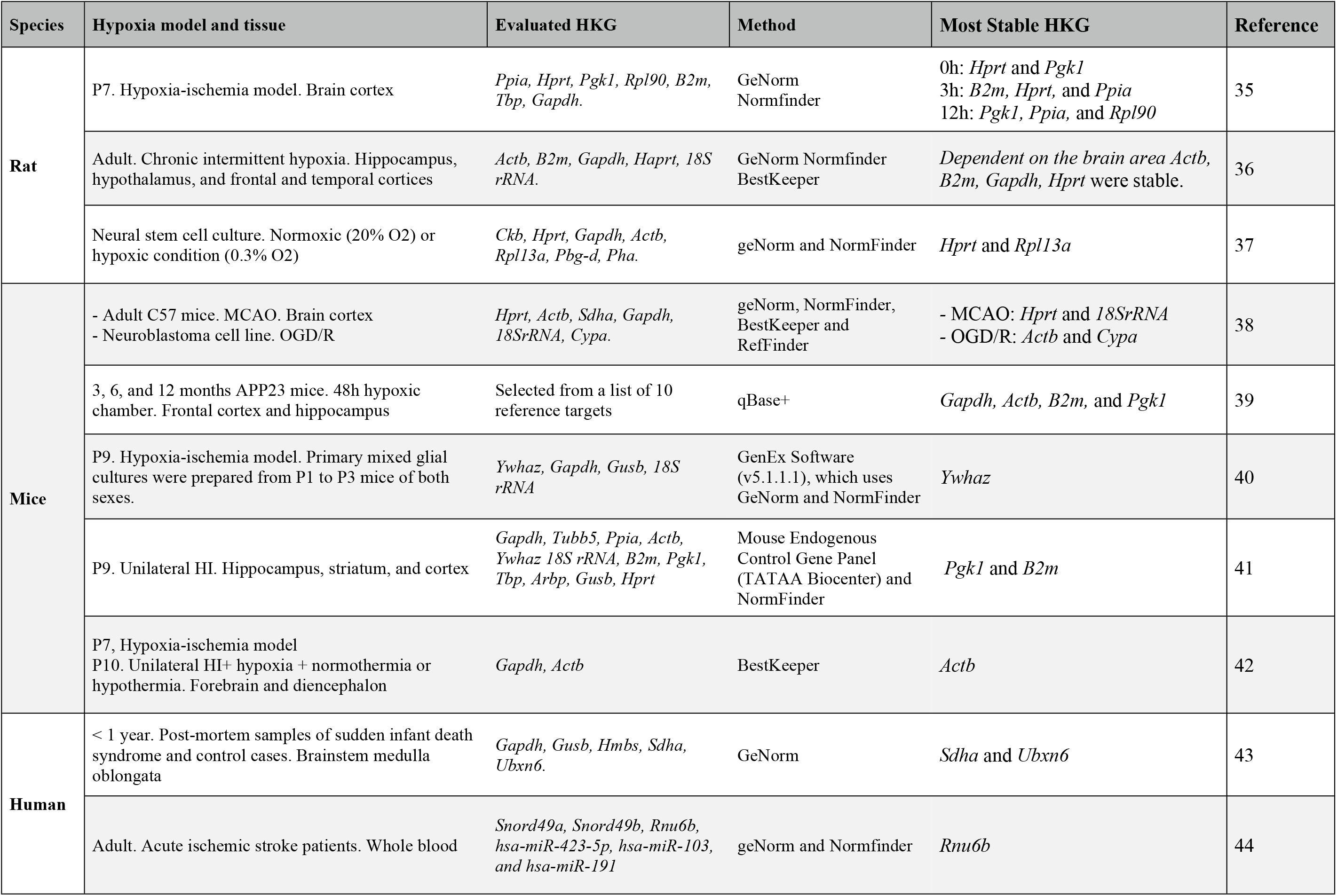
List of published RG validation studies involving hypoxia.

## Methods

### Ethics statement

This study was carried out in accordance with the recommendations in the Guide for the Care and Use of Laboratory Animals of the National Institute of Health of Argentina. The protocol was approved by the Biomedical Ethics Committee of Universidad Católica de Cuyo, San Juan, Argentina and by the Ethical Committee of CICUAL: “Institutional Committee for the Use and Care of Laboratory Animals” (Resolution no. 2079/07), Facultad de Medicina, Universidad de Buenos Aires, Argentina. Appropriate actions were taken to minimize the number of animals used and their suffering, pain, and discomfort.

### Hypoxic-ischemic injury animal model

Severe acute PA was induced using a model of hypoxia-ischemia as described previously (26–28). Briefly, albino Sprague-Dawley rats were kept under standard laboratory conditions at 24°C with light-dark cycles of 12:12 hours, and food and water present ad libitum. Fifteen timed-pregnant Sprague-Dawley rats were used. The first group of offspring studied consisted of normally delivered naive pups that were used as controls (CTL; n=6). After vaginal delivery of the first pup, pregnant dams were euthanized by decapitation and immediately hysterectomized. All full-term fetuses, still inside the uterus, were subjected to asphyxia by transient immersion of both uterine horns in a saline bath for 20 min at either 37°C (perinatal asphyxia in normothermia, PA, n=6) or 15°C (perinatal asphyxia in hypothermia, [HYPPA]; n=6). After asphyxia, the uterine horns were opened, pups were removed, dried of delivery fluids, and stimulated to breathe, and their umbilical cords were ligated. After recovery, one group of PA animals was placed on a cooling pad at 8ºC for 15 minutes for hypothermic treatment (PAHYP, n=6), while hypothermic control animals (HYP, n=6) received the same treatment. After 15 minutes of exposure to the cold environment the core temperature of the newborns was measured with a rectal probe (mean temperature: 20.1ºC; n=8). The pups were subsequently placed under a heating lamp for recovery after which they were and placed with a surrogate mother. Time of asphyxia was measured as the time elapsed from the hysterectomy up to the recovery from the water bath. Pups that adjusted to the following parameters were included: 1. Occipitocaudal length > 41mm, 2. Weight > 5g.

### Total RNA extraction and reverse transcription cDNA synthesis

Animals were sacrificed by quick decapitation 24 h post treatment. The brain cortex was isolated, snap-frozen in liquid nitrogen, ground into powder with pestle and mortar cooled in liquid nitrogen and then stored at −80 °C. Total RNA was isolated from about 80 mg tissue powder using TRIzol® (Invitrogen Life Technologies, USA) following the manufacturer’s instructions. The residual DNA was removed by the TURBO DNA free kit (Ambion Inc., UK). Yield and purity of RNA was determined by the NanoDrop ND-1000 spectrophotometer (Nanodrop technologies, USA). RNA samples with an absorbance ratio OD 260/280 between 1.9–2.2 and OD 260/230 greater than 2.0 were used for further analysis. RNA integrity was assessed using agarose gel electrophoresis. One microgram of RNA from each sample was reverse transcribed using the High Capacity cDNA Reverse Transcription Kit (Applied Biosystems) according to the manufacturer’s instructions. cDNA was stored at −20 °C for future use. For qPCR analysis, each cDNA sample was diluted 20 times with nuclease-free water.

### Real-time PCR

Real-time PCRs were conducted using the LightCycler® 480 Multiwell Plate 96 (Roche, Mannheim, Germany) containing 1µM of each primer. For each reaction, the 20 μl mixture contained 1 μl of diluted cDNA, 5 pmol each of the forward and reverse primers, and 10 μl 2 × SensiMix SYBR No-ROX Kit (Bioline, UK). The amplification program was as follows: 95°C for 30 sec, 40 cycles at 95°C for 15 sec, and 60°C for 15 sec, and 72°C for 15 sec. After amplification, a thermal denaturing cycle was conducted to derive the dissociation curve of the PCR product to verify amplification specificity. Reactions for each sample were carried out in triplicate. qPCR efficiencies in the exponential phase were calculated for each primer pair using standard curves (5 ten-fold serial dilutions of pooled cDNA that included equal amounts from the samples set). The mean threshold cycle (Ct) values for each serial dilution were plotted against the logarithm of the cDNA dilution factor and calculated according to the equation E = 10(−1/slope) × 100, where the slope is the gradient of the linear regression line.

### Reference gene selection

Based on their common usage as endogenous control genes in previous studies (Table 1), five candidate RGs were analyzed, i.e., *Actb*, *Pgk1*, *Gapdh*, *Sdha*, *Rnu6b*. These genes represent commonly used endogenous control genes chosen from the relevant literature and have been previously validated in rat, mouse and human brain tissues exposed to hypoxia. The selected RGs belong to different molecular pathways to minimize the risk of co-regulation between genes. The primers were designed from nucleotide sequences identified using NCBI BLAST (http://blast.ncbi.nlm.nih.gov/Blast.cgi). *Rnu6b* TaqMan MicroRNA Assay (*Rnu6b*) was commercially available (Thermo Fisher Scientific, Product number: 4427975-001093). All other primers were ordered from Thermo Fisher Scientific with their certificates of analysis. The primer characteristics of nominated RGs are listed in Table 2. The primer sequences (5’-3’) of the target genes were as follows:

*Rest*; - F, AACTCACACAGGAGAACGCC - R, GAGGTTTAGGCCCGTTGTGA.

*Bad*; - F, GCCCTAGGCTTGAGGAAGTC - R, CAAACTCTGGGATCTGGAACA.

**Table 2.**
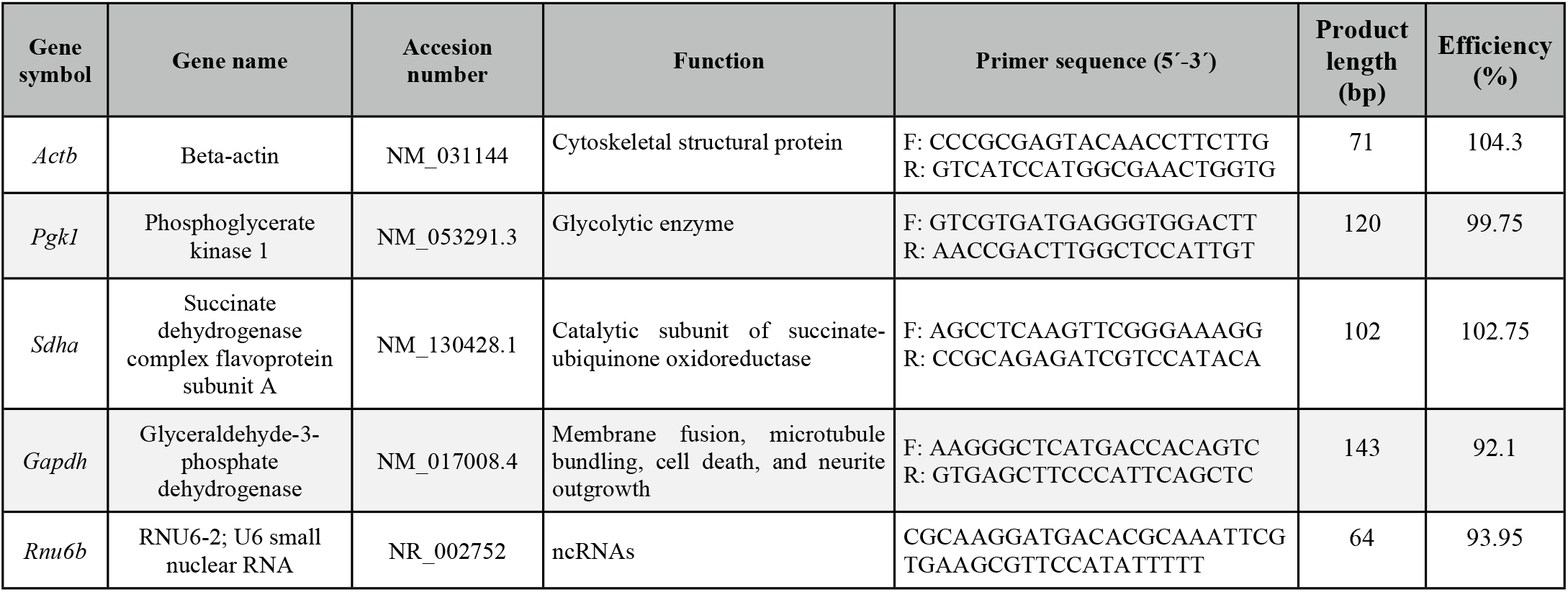
List of RGs investigated by qPCR.

#### Analysis of expression stability using multiple statistical approaches

To assess the stability of candidate RGs, five statistical methods, each with unique characteristics, were used: GeNorm, BestKeeper, NormFinder, Coefficient of Variation analysis, and the comparative ΔCt method. Ct values were converted to non-normalized relative quantities according to the formula: 2−ΔCt. CV analysis, GeNorm and NormFinder calculations are based on these converted quantities, whereas BestKeeper and the ΔCt method make use of raw Cq values.

#### Impact of selection of RGs on gene expression normalization

The impact of RG selection on gene expression quantification was assessed via examining the expression of *Rest* and *Bad.* Six gene expression normalizing strategies were used to represent the least and most stable reference genes. The relative expression profiles of *Rest* and *Bad* were determined and normalized with all tested RGs. Relative fold changes in gene expression were calculated using the DDCt and Pfaffl methods. Data was expressed as mean ± standard error of the mean (SEM) from six independent samples/group with triple qPCR reactions. One-way analysis of variance (ANOVA) test was applied to analyze significant differences between conditions for each house-keeping gene. Differences were reported as statistically significant when *p*<0.05. GraphPad Prism 6 (GraphPad Software, USA) was used for statistical procedures and graph plotting.

## Results

### qPCR

Pilot assays were performed to optimize cDNA and primer quantities. A total of 0.9 mg of RNA that was previously treated with DNase was used for the reverse transcription reaction in a total volume of 40ml. One microliter of the resulting cDNA was used for the qPCR reaction. Each gene amplification was analyzed, and a melting curve analysis was performed, showing a single peak indicating the temperature of dissociation. Efficiencies are shown in Table 2. All Ct values were between 17.0 and 33.0.

### Coefficient of Variation analysis

We calculated the raw expression profiles of RGs as changes of Ct values across groups and ranked the gene stability according to the CV. The CV analysis is a descriptive statistical method where the Ct values of all candidate RGs across samples are first linearized (2^−Cq^). Next, the CV for each gene across all samples was calculated and expressed as a percentage. The CV estimates the variation of a gene across all samples taken together, and, therefore, a lower CV value indicates higher stability. This analysis on the cortical samples revealed *Gapdh* as the most stable RG, and *Actb* as the least stable RG. This method however does not consider the variation across different treated groups; hence, CV analysis alone cannot determine the best set of RGs.

To assess if the mean mRNA levels across groups were significantly different from one another, a One-way ANOVA was used. The results demonstrated that variations in the Ct values for the different treatments were different for all candidate RGs. Four of the five genes tested (*Sdha, Rnu6b, Pgk1, Actb*) showed significant variation in mRNA levels across different treatments (Fig 2). Only *Gapdh* showed no significant changes. These results, making use of the raw expression profiles of the RGs, suggest that the various experimental conditions were associated with changes in RG expression levels that, as such, could skew the normalized profile of target genes. As a result, RG selection without accounting for potential expression differences between conditions is accompanied by a significant bias in the results and their interpretation. Hence, it is of utmost importance to validate the stability of RGs prior to normalization in gene expression studies.

**Fig 1.**
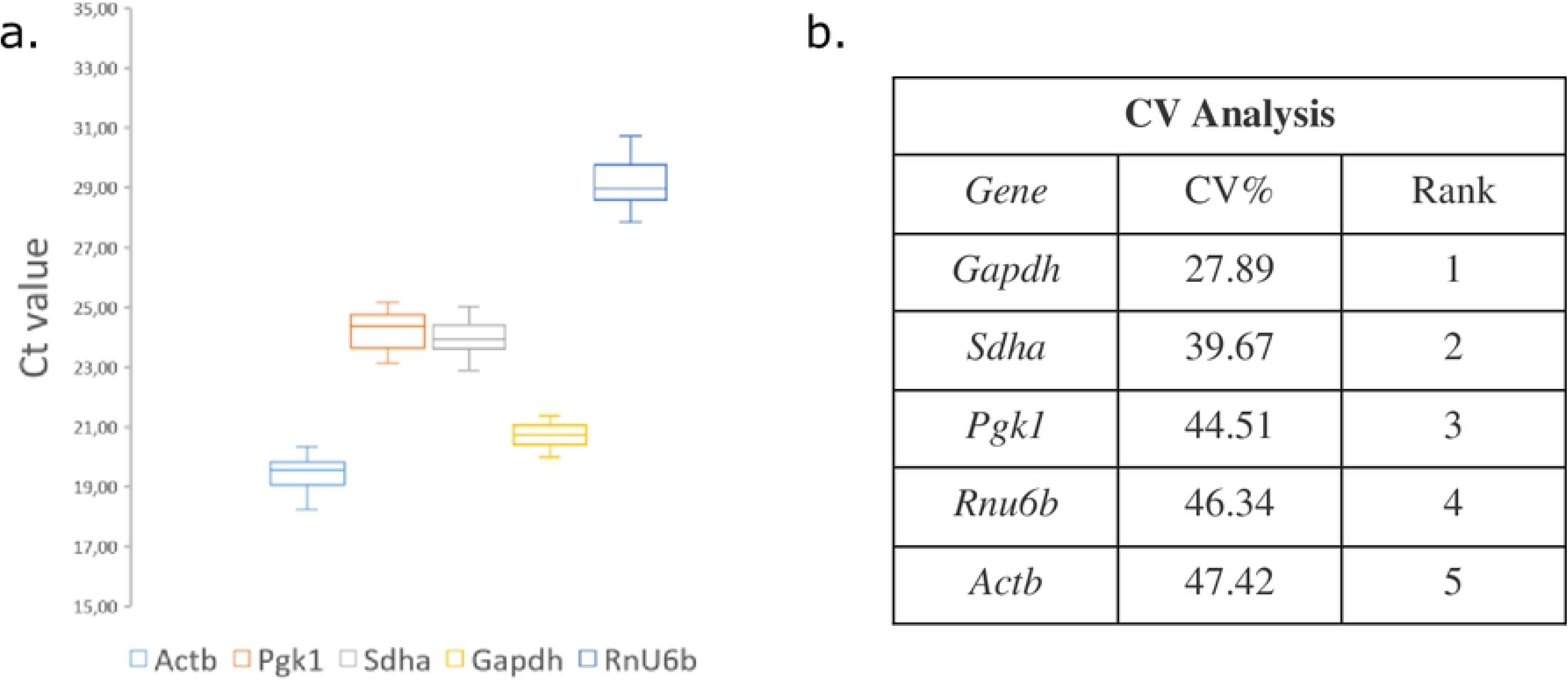
Variability of the raw Ct values of the five candidate RGs under different experimental conditions. a. Relative quantities without normalization to any RG using cerebral cortex samples (n=30). The boxes encompass the 25th to 75th percentiles, whereas the line in the box represents the mean. Whisker caps denote the maximum and minimum values. b. CV analysis of the linearized Ct values.

**Fig 2.**
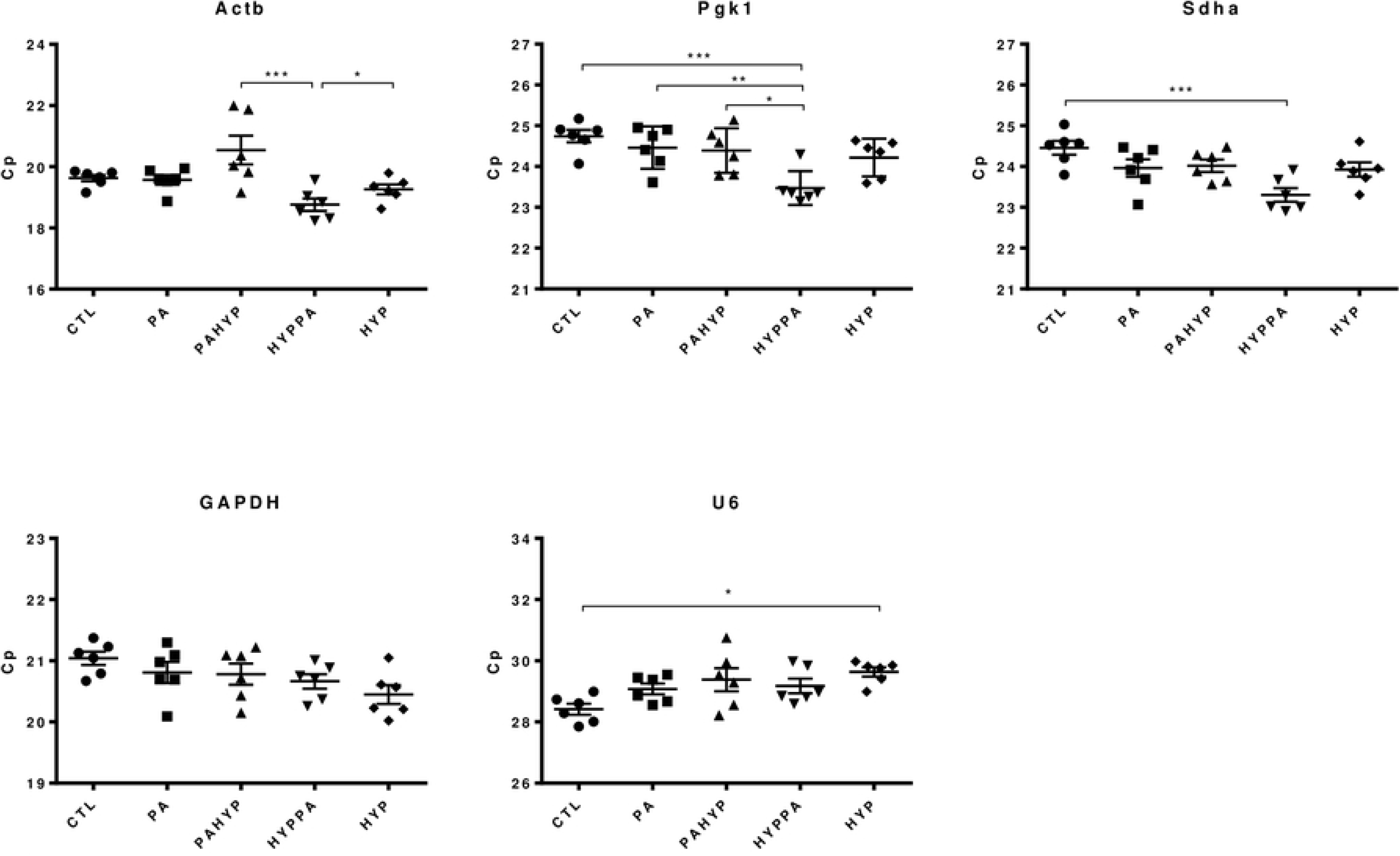
Expression profiles of RG expressed as Cp across the experimental conditions. a. *Actb*, b. *Pgk1*, c. *Sdha*. d. *Gapdh*, e. *Rnu6b.* Results are expressed as the Mean ± SEM for each treatment. One-way ANOVA was performed to asses differences between the means of all groups. Statistical significance is denoted by *p* values: \**p*<0.05, \*\**p*<0.01, \*\*\**p*<0.001.

Next, to identify the optimal RG(s), the expression stability of candidate RGs was analyzed using four well known statistical methods (Table 3).

**Table 3.** Candidate RG expression stability.

| Rank | GeNorm |  | NormFinder |  | BestKeeper |  |  | Δ Ct method |  | Comprehensive ranking |  |  |  |
| --- | --- | --- | --- | --- | --- | --- | --- | --- | --- | --- | --- | --- | --- |
|  | Gene | M | Gene | S | Gene | Cv<br>(%Ct) | SD<br>(±Ct) | r | Gene | Mean<br>SD | Geomean | Rank | Gene |
| 1 | *Pgk1* | 0.596 | *Actb* | 0.222 | *Pgk1* | 2.17 | 0.53 | 0.825 | *Pgk1* | 1.41 | 1.5 | 1 | *Pgk1* |
| 2 | *Actb* | 0.599 | *Sdha* | 0.298 | *Actb* | 2.83 | 0.55 | 0.819 | *Sdha* | 1.45 | 2 | 2 | *Actb* |
| 3 | *Sdha* | 0.782 | *Pgk1* | 0.298 | *Sdha* | 1.54 | 0.32 | 0.814 | *Actb* | 1.53 | 2.5 | 3 | *Sdha* |
| 4 | *Gapdh* | 1.053 | *Gapdh* | 1.736 | *Gapdh* | 1.84 | 0.44 | 0.614 | *Gapdh* | 2.00 | 4 | 4 | *Gapdh* |
| 5 | *Rnu6b* | 1.923 | *Rnu6b* | 3.17 | *Rnu6b* | 1.97 | 0.57 | 0.106 | *Rnu6b* | 3.23 | 5 | 5 | *Rnu6b* |

Stability was ranked by GeNorm, NormFinder, BestKeeper and $\Delta$ CT average STDEV. The comprehensive ranking was based on the geometric mean of the gene rank. Candidates are listed from top to bottom in order of decreasing expression stability. (SD [ ± Ct]: standard deviation of the Ct; CV [% Ct]: coefficient of variance expressed as a percentage of the Ct level; geomean: geometrical mean).

First, a GeNorm analysis was performed on all five candidate genes. GeNorm calculates stability value (M) based on pairwise variation of every two genes. The rationale is that if two genes vary similarly across all samples, then they are the most stable RGs for that dataset. A limitation of this method is that if two genes are regulated in the same direction by one or more experimental conditions, those will often be assumed to be the most stable. In our analysis, except for *Rnu6b*, which presented the highest M-value (M=1.923), all of the other candidate RGs presented M-values lower than 1.5, which is considered to be the cut-off for suitability. Based on this analysis for the neonatal cortex, the most stable RGs were *Pgk1* and *Actb*. This is in stark contrast to the CV analysis, that showed those genes as the least stable ones (higher CV), and to the expression profiles that showed inter-group differences.

NormFinder calculates the stability score (S) based on the inter- and intra-group variation. However, it has been reported that including genes with high overall variation can affect the stability ranking of all genes with this method (22). This algorithm can potentially be improved after identifying and removing genes with high overall variation.. *Actb*, *Sdha* and *Pgk1* were the most stable RGs, presented stability values lower than 0.3. *Gapdh* (SV=1.736) and *Rnu6b* (SV=3.17) were the least stable.

BestKeeper uses the cycle threshold (Ct) values to calculate a standard deviation (SD), coefficient of variance (CV), and Pearson correlation coefficient (r) for each gene. Lower SD and CV values indicate more stable gene expression, and genes that exhibit a SD in Ct values above 1.0 should be eliminated and regarded as unreliable controls. Then, the remaining RG are ranked according to r values, with a higher r value indicating more stable gene expression. None of the genes analyzed were excluded for having SD above 1. The most stable RG was *Pgk1* (r=0.825), while *Rnu6b* was considered the least stable gene (r=0.106). The ranking obtained from this analysis was the same as the one obtained with GeNorm.

The Δ-Ct method works on the same rationale as GeNorm but calculates the stability value (mean SD) differently; it is calculated as the average standard deviation of the Ct value differences that the gene exhibits with other genes. Using this method, the ranking was similar to previous rankings. The most stable RGs were *Pgk1* (Av. SD=1.41) and *Sdha* (Av.SD=1.45), and the least stable *Rnu6b* (Av.SD=3.23). The overall ranking depicted in Table 3 was based on the geometric mean of the previous gene ranks. This ranking indicates that for this tissue and treatment, the most stable RG was *Pgk1*.

### Impact of RG selection on target gene expression profiles

The impact of RG selection on gene expression quantification was assessed by examining the expression of *Rest* and *Bad*. These genes have shown to be influenced by hypoxia and hypothermia. Five gene expression normalizing strategies were used to select the least and most stable RGs, and the best combination of two genes, *Actb/Pgk1* (Fig 3). Expression values were calculated relative to expression in control animals, using both the **ΔΔ**Ct method (Livak & Schmittgen, 2001) and the primer efficiency method (Pfaffl, 2001, Fig 3). Results were similar using Livak or Pfaffl methods. As expected, even when the general pattern of target gene expression was similar for most of the RGs across treatments, target gene expression levels were different depending on the RG used for normalization causing differences in the significance level of the expression patterns.

**Fig 3.**
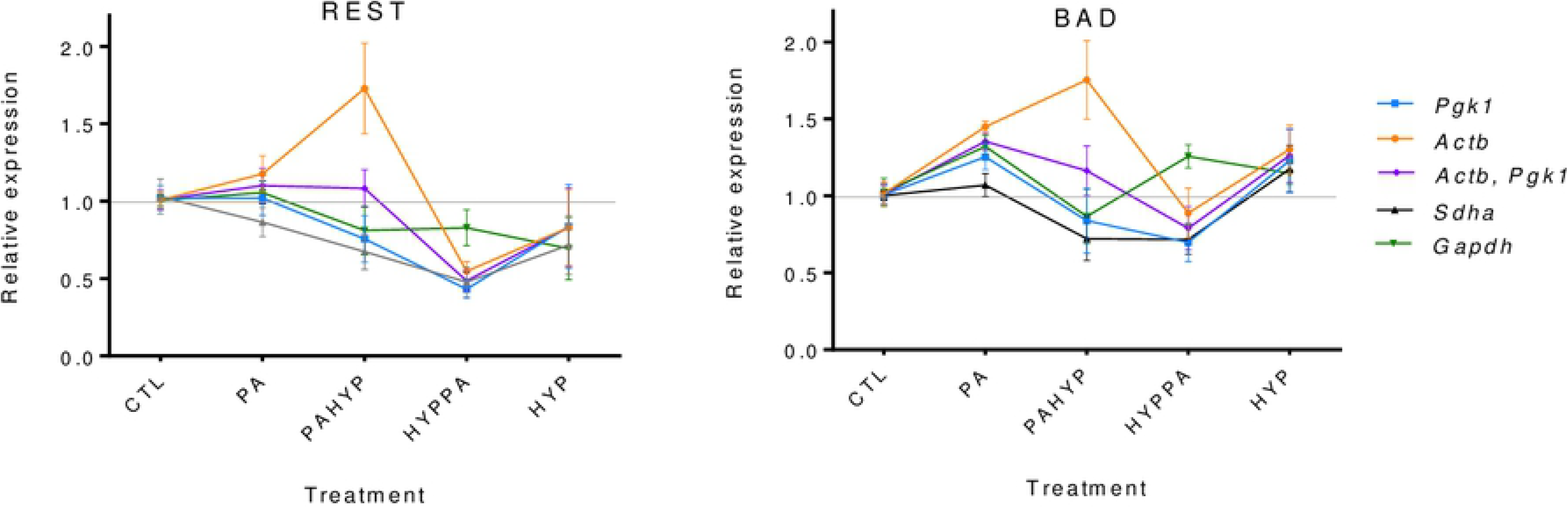
Evaluation of the impact of selection of RG on gene expression normalization. Expression profiles of *Rest* and *Bad* normalized by different strategies. Arithmetic mean values and standard deviations were obtained from three bioreplicates.

## Discussion

The selection of RGs in qPCR experiments has an enormous impact on the reliability and interpretation of results in gene expression studies making it a crucial yet often understated process. It is now recognized that normalization of qPCR results against a single RG is likely to be inadequate and that normalization against a panel of RGs containing at least three stable RGs is preferred. However, for most of the RGs used in published qPCR studies, no thorough investigation of their variation over experimental conditions has been performed and/or reported (48). Many researchers continue to use a single, unvalidated RG to normalize data.

The majority of studies where assessment of the RGs’ stability is included make use of statistical tools like GeNorm, BestKeeper, NormFinder, CV analysis, and the comparative ΔCt method. Each of these methods determines the stability based on a set of assumptions and calculations, and has its own limitations. In general, methods that rely on pairwise variation (GeNorm and Pairwise ΔCt method) are influenced by the expression pattern of all genes making their ranking inter-dependent. The CV analysis does not take the variation between groups into account, hence alone it cannot determine the best (set of) RG(s), but it can be used as a first filter to discard genes with high overall variance. Moreover, except for the CV analysis, the presence of genes with high overall variation impact upon the ranking of all these methods.

As a result, the selection of stable RGs varies significantly depending on the method used making the choice of the validation method a critical step in qPCR assays. In our study, using Geomean, *Pgk1* was the most stable gene across treatments, while *U6* and *Gapdh* were ranked as most variable. This is in stark contrast to the CV% Analysis and intergroup ANOVA Ct variations that indicated that *Gapdh* was the most stable gene among groups, and *Actb* the least stable.

Using any of these methods alone is not sufficient in obtaining bias-free results. Generally, stability validation studies have ranked the genes using Geomean, a ranking obtained from the mean rank of the statistical tools used. This method does not take into account the limitations of each algorithm separately, which is why it is increasingly considered an erroneous approach when validating RGs. This makes the identification of the best RGs very unwieldy. Using the same statistical methods, new approaches have been proposed, such as the “Integrated approach” (22) that has shown to provide a more accurate estimate of RG stability. It is advisable to devise integrated approaches based on suitability for each experimental setting.

Although we analyzed a small set of candidate RGs, we found differences in the stability rankings obtained with the different methodologies, and the associated bias was reflected in our target gene quantification. Our study emphasizes the necessity of validating RGs previous to assessing target gene qPCR data, and the importance of choosing the right set of statistical methods for doing so. Such an approach would lead to more accurate and reproducible expression assessment.

## Abbreviations

qPCR: real-time reverse transcription
RGP: reference gene
*Actb*: beta-actin
*Pgk1*: phosphoglycerate kinase 1
*Sdha*: succinate dehydrogenase complex flavoprotein subunit A
*Gapdh*: glyceraldehyde-3-phosphate dehydrogenase
*Rnu6b*: U6 small nuclear RNA
18S rRNA: 18S ribosomal RNA
*Hpr*: hypoxanthine-guanine phosphoribosyltransferase
*B2m*: beta-2-micro-globulin
*Tubb5*: tubulin beta 5
*Ppia*: peptidylprolyl isomerase A
*Ywhaz*: tyrosine 3/tryptophan 5-monooxygenase activation protein zeta polypeptide
*Pgk1*: phosphoglycerate kinase 1
*Tbp*: TATAA-box binding protein
*Arbp*: acidic ribosomal phosphoprotein P0
*Gusb*: beta-glucuronidase
*Ckb*: brain creatine kinase
*Rpl13a*: ribosomal protein L13A
*Pbg-d*: porphobilinogen deaminase
*Cypa*: cyclophilin
*Rest*: repressor element 1-silencing transcription factor
*Bad*: BCL2/BCL-XL-associated death promoter

